# Improved Peptide Docking with Privileged Knowledge Distillation using Deep Learning

**DOI:** 10.1101/2023.12.01.569671

**Authors:** Zicong Zhang, Jacob Verburgt, Yuki Kagaya, Charles Christoffer, Daisuke Kihara

## Abstract

Protein-peptide interactions play a key role in biological processes. Understanding the interactions that occur within a receptor-peptide complex can help in discovering and altering their biological functions. Various computational methods for modeling the structures of receptor-peptide complexes have been developed. Recently, accurate structure prediction enabled by deep learning methods has significantly advanced the field of structural biology. AlphaFold (AF) is among the top-performing structure prediction methods and has highly accurate structure modeling performance on single-chain targets. Shortly after the release of AlphaFold, AlphaFold-Multimer (AFM) was developed in a similar fashion as AF for prediction of protein complex structures. AFM has achieved competitive performance in modeling protein-peptide interactions compared to previous computational methods; however, still further improvement is needed. Here, we present DistPepFold, which improves protein-peptide complex docking using an AFM-based architecture through a privileged knowledge distillation approach. DistPepFold leverages a teacher model that uses native interaction information during training and transfers its knowledge to a student model through a teacher-student distillation process. We evaluated DistPepFold’s docking performance on two protein-peptide complex datasets and showed that DistPepFold outperforms AFM. Furthermore, we demonstrate that the student model was able to learn from the teacher model to make structural improvements based on AFM predictions.

## 1. Introduction

Protein-peptide interactions are crucial in many biological processes and are frequently used in the early stages of many drug development pipelines, as they can often help better understand how to target a protein of interest. Furthermore, many proteins have highly flexible peptide-like intrinsically disordered regions that are crucial for biological functions and can mediate up to 40% of all protein interactions [1]. However, experimental determination of protein-peptide complexes via methods such as X-ray crystallography, cryo-electron microscopy, and nuclear magnetic resonance spectroscopy are slow and resource intensive, thus there are limited structures available for protein-peptide complexes in the Protein Data Bank (PDB) [2]. Consequently, there has been a growing effort to develop in silico methods for the structure modeling of protein-peptide complexes, as it can provide significant insight without the time or expense experimental methods require. Computational methods have been developed for a wide range of tasks related to protein-peptide interactions, ranging from predicting peptide binding residues [3] to directly predicting protein-peptide complexes [4]. Nonetheless, due to the flexible nature of peptide structures, predicting protein-peptide complexes remains a challenging problem in comparison to modeling protein-protein complexes. Recent advances in deep learning have led to increased efforts in developing structure prediction tools using deep neural networks [5], [6]. However, most methods have focused on predicting protein-peptide interactions or binding residues rather than modeling the peptide-receptor complex. AlphaFold (AF) has significantly elevated the field of structural biology, allowing for the direct prediction of structures from sequences in an end-to-end fashion with high accuracy [7]. Even though AF is primarily trained on monomeric structures, it has been shown that AF can predict complex structures with minor modifications, such as adding residue gaps between chains [8] or using a linker [9]. With the release of AlphaFold-Multimer (AFM) [10], a re-trained version of AF using multimeric structures, it is now possible to model protein complexes with more accurate interfaces compared to previous methods that relied on modifying AF inputs. Several studies have also evaluated the performance of AFM in modeling protein-peptide complexes [11], [12]. These studies have shown that there are often substantial errors in modeling of the protein-peptide complexes, particularly in targets with sufficiently long peptide sequences indicating that there is still room for improvement in accurately modeling protein-peptide complexes.

Knowledge Distillation (KD) is a well-established concept in the field of machine learning, offering a mechanism for transferring knowledge from one neural network to another. This concept draws inspiration from the way humans learn, employing a framework akin to a teacher guiding a student during the learning process. In this paradigm, a proficient teacher network imparts its knowledge to a less proficient student network through a training process. KD has proven to be particularly useful in enhancing the learning capabilities of the student network. There are two main categories within the realm of KD:

1. Model-based distillation: In this approach, a teacher network with high model complexity shares its knowledge with a student network designed to be much smaller and computationally efficient [13], [14]. This process helps the student network learn important features and relationships without being burdened by excessive complexity.
2. Feature-based distillation, which is also referred to as privileged knowledge distillation (PKD): In this context, the teacher network has privileged access to additional information during training, and it strives to impart this privileged knowledge to the student network, which does not possess the same access [15]. PKD has shown success in a variety of computer vision tasks, including image segmentation [16] and object detection [17].

Building upon the promising developments in KD, here we introduce a novel approach, DistPepFold (Peptide docking using Distillation) for the modeling of complexes between globular proteins and peptides or other flexible or disordered protein regions. DistPepFold leverages PKD to enhance the structural modeling of protein-peptide complexes using AFM. DistPepFold consists of two integral components: 1. Teacher model: This model utilizes native interaction information as privileged information during training. 2. Student model: The student network learns from the teacher, absorbing the valuable knowledge and guidance imparted by the teacher. Both the teacher and student models use AFM’s single and pair representations as input and directly predict the 3D coordinates of the protein-peptide complex structure. Our approach employs the teacher’s predicted structure and intermediate representations to guide the learning process of the student network. Through rigorous evaluation on two datasets of protein-peptide complexes, we demonstrate the efficacy of our proposed method. Notably, the student model consistently outperforms AFM using traditional docking analysis metrics, showcasing the benefits of knowledge distillation in this context. Moreover, in cases where AFM has low confidence scores on its predicted structures, our method improved the modeling of protein-peptide interactions compared to the AFM predicted structure.

## 2. Methods

### 2.1 Overall framework

Here we present the details of DistPepFold (**Figure 1**). The goal of the proposed framework is to train a student model who can improve predicted structures from AFM. This is achieved by making a student model learn better single and pair representations from a well-trained teacher model that uses native-interaction information. Given sequences of the peptide and receptor as input, we first run AFM to generate structures and intermediate representations. Here, the intermediate representations consist of the single representation (L x 384), and pair representation (L x L x 128), where L denotes the length of the input sequence. To extract the representations from AFM, we used the single and pair representations from the last recycle iteration. Next, we train the teacher model with native interaction information, where the teacher model learns from both AFM representations and native structures. The teacher model consists of three components: 1) the contact encoder (CE), 2) the trunk and 3) the structure module (SM). Then, after the teacher model has completed training, we use the teacher models’ representations and predicted structure as hint knowledge to train a student model of similar architecture on the same training set. During inference, we first run AFM and then use the representations as input to the student model. The student model will directly output the 3D coordinates of the structures.

**Figure 1:**
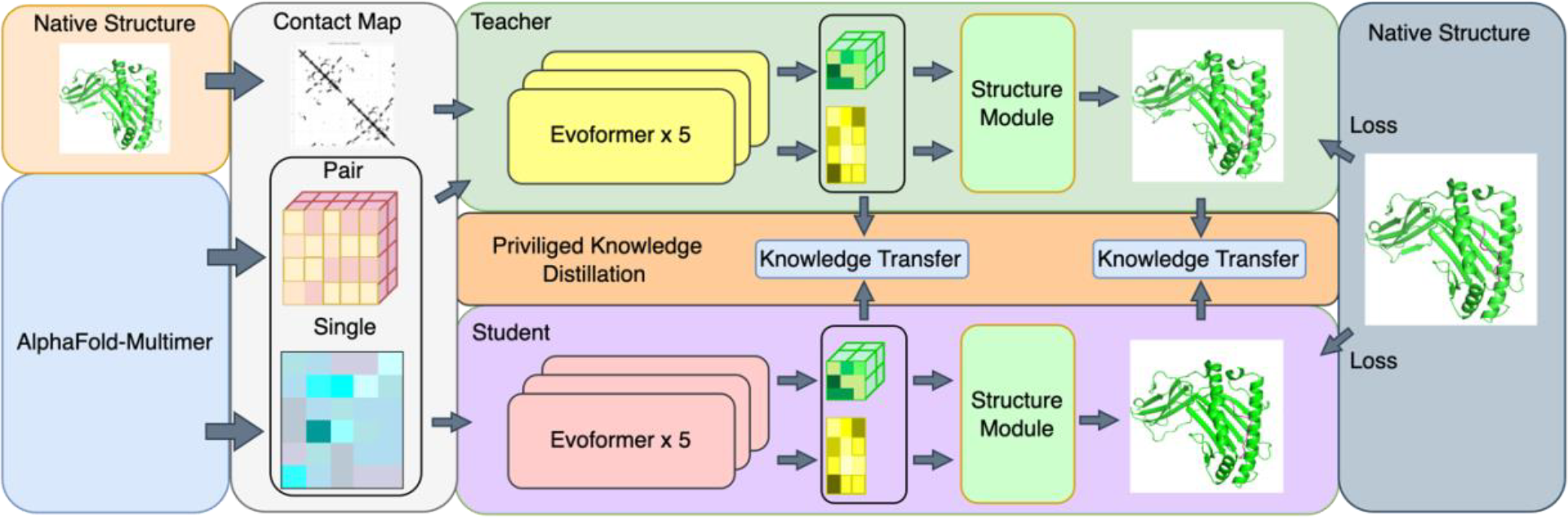
Overall framework of DistPepFold. DistPepFold consists of a teacher model and a student model. The input to the teacher model includes single and pair representations, as well as the contact map over the interaction region from the native structure. The input to the student model includes only the single and pair representations. Both single and pair representations are generated from AFM. The teacher and student models share similar architecture and largely follow that of AFM, each containing 5 blocks of Evoformers and a structure module.

### 2.2 Network Architecture

Our proposed method has two networks: a teacher model and a student model. The teacher model is a native-interaction-privileged structure prediction network. The model consists of three parts: 1) the contact encoder (CE) that encodes contact information. 2) the trunk that uses Evoformer architecture from AFM. 3) the structure module (SM) that predicts 3D structure coordinates from the single representation. The teacher model uses single and pair representations as input. In addition, we added contact information over the interaction region as input to the teacher model. We first converted the coordinates of the native structures into binary contacts, in which two residues are considered to be in contact if their Cα-Cα distance is within 8 Å. The processed contact map is input into CE to generate high-level features to be used in later parts of the model. CE leverages two blocks of ResNet [18]. Each ResNet block consists of two 2D-convolution layers with 128 channels, a kernel size of 3 and a skip connection. The output of CE will be fused with pair representations before feeding to the trunk. To fuse the information, we simply performed element-wise addition of the representations. The trunk consists of 5 blocks of Evoformer. Since we only use the single and pair representations from AFM, we eliminate column-wise attention from the original Evoformer implementation. The rest of the architecture remains the same as the proposed Evoformer used in AF. Lastly, the trunk will output the processed single and pair representations and feed them into the SM for structure prediction. The student model shares the same architecture as the teacher model, with the exception of the CE, thus containing only 5 blocks of Evoformer and a structure module. The student model directly takes single and pair representations as input and predicts the coordinates of the structure.

### 2.3 Loss Function

To train the teacher model, we used the structure loss as proposed in AF [7]. The structure loss consists of the Frame Aligned Point Error (FAPE) loss and a series of auxiliary losses. The structure loss for the teacher model can be defined as follows:

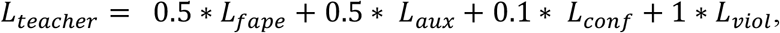

where *L*_*fape*_ and *L*_*aux*_ are FAPE computed over sidechain atoms and mainchain Cα atoms respectively. In addition, *L*_*aux*_ includes torsion angle losses as defined in AF. *L*_*conf*_ is a confidence score computed to estimate model accuracy in the form of predicted TM-score (pTM), which can be directly computed from the pair representation during training [10]. During inference, the confidence of predicted structures is a weighted combination of pTM score computed over interaction and non-interaction regions. *L*_*viol*_ is the structure violation loss that penalizes any atom clashes and encourages an acceptable quality of the predicted structures. We did not use distogram loss and MSA loss from AF since they do not apply in our framework. We also omit the experimentally resolved loss term as we did not observe improvements with the loss term. Similar to AFM, backbone FAPE over interaction regions is up-weighted by using a clamping value of 30 compared to a clamping value of 10 for the non-interaction regions. In addition, to force the model to pay more attention to the peptide structure, we up-weighted the FAPE over the peptide region using a clamping value of 30 as well. We adopt the same weight value associated to each loss term as in AF, since we observed no significant improvements through tuning these hyperparameters.

To train the student model, we follow the PKD strategy described previously [19]–[21]. First, we define the structure loss for student as *L*_*structure*_, which is computed between the student model’s predicted structure and the native structure. We used the same formula as *L*_*teacher*_ as described above. In addition to the structure loss, we aim to transfer the knowledge from the teacher model to the student model through soft label loss and latent space loss. We define soft label loss *L*_*soft*_ as the structural loss between the student’s predicted structure and the teacher’s predicted structure, which can be described as:

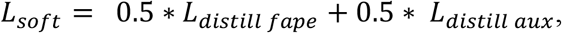

where *L*_*distill fape*_ and *L*_*distill aux*_ are FAPE compute for sidechain atoms and mainchain Cα between the student and teacher’s structure outputs respectively. We omit the violation term due to training stability issues. The idea of using soft label loss is to discover the relation between representations and structures that are hard to optimize during training, which has been shown to improve performance by previous knowledge distillation studies [22]. Moreover, we define latent space loss *L*_*latent*_ as the loss between the student and teacher’s single and pair representations, which can be described as:

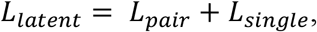

where *L*_*pair*_ is mean absolute error loss (L1 loss) between student and teacher’s pair representations, which contain residue-residue distance information. Pair representations from the teacher model contain native interaction information, especially over the interaction region. This loss encourages the student to learn from such information and mimic the representation from the teacher when native information is not presented. *L*_*single*_ is the L1 loss between the student and teacher’s single representations. The single representation contains per-residue level information, which yields the final coordinates of the structure. The idea of using *L*_*latent*_ is to enforce the student to not only produce teacher-like representations at the last layer of the network, but to also produce teacher-like representations at intermediate layers of the network. This is often referred to as “hint learning” in the general knowledge distillation framework [22], [23]. Both *L*_*soft*_ and *L*_*latent*_ are considered as distillation loss in our framework to ensure that the student model can learn useful information for the teacher model. The total loss function for student *L*_*student*_ consists of *L*_*structure*_, *L*_*soft*_ and *L*_*latent*_, which can be described as:

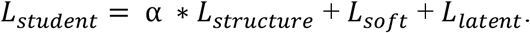

We observed that FAPE between the predicted structure and the ground truth has strong regularization effects in which it often leads training optimization into a certain local minima. It is possible that some structures are hard to learn from the ground truth but are easier to learn from the teacher’s model. Thus, we use hyperparameter α to control this regularization effect. Lower values of α indicates that less constraints are applied to the students to learn from the ground truth and encourages the student to learn more from the teacher model.

## 3 Experiment

### 3.1 Dataset

We used pre-formatted files from the 2020-03-18 release of the PepBDB [24] to collect protein-peptide complexes. PepBDB is a publicly available database that is designed for studying protein-peptide interactions. Since we are specifically interested in longer peptides that are ordered when bound to a receptor surface, we applied several filters to remove shorter peptides, and those that lack a globular receptor. From the 13,527 entries in the PepBDB, we filtered out entries which 1) have resolution lower than 3 Å. 2) have a peptide length less than 10 resolved residues. 3) have a receptor structure with less than 100 resolved residues. We also removed cases in which the same PDB ID was listed in multiple PepBDB entries. This resulted in 3,975 entries. We then used MMseq2 [25] to cluster the sequences with 40% sequence similarity using the receptor sequence, which resulted in 729 sequence clusters. We separated the 729 clusters into training, validation, and testing with a ratio of 7:1:2. During training, we considered all redundant sequences. This resulted in 3,048 training entries, 45 validation entries and 92 testing entries. From this data, we created cleaned and renumbered Fasta and PDB files, and generated embeddings with the AlphaFold2 v2.2.0 release [26].

To increase the amount of training data available, we derived peptide-protein complexes from globular complexes present in the 2021-01-05 release of the PDB, as globular complex structures have previously proven useful as peptide templates [27]. From these complexes, we ran Rosetta Peptiderive [28], [29] on all pairwise combinations of chains within all PDB entries. Peptiderive works by using a sliding window of amino acids (here we used 15 amino acids) of one chain, while keeping the other fixed. The window with the best binding energy is kept and exported as a potential peptide. From all generated peptide-protein complexes, we remove cases in which less than 60% of the peptide is within 5 Å of the receptor, where there were less than 150 residues within the receptor, and cases in which cyclic peptides were generated. Although the complexes are not real peptide-protein complexes, the interactions between the fragment and receptor structures can still be useful in training. We noticed that AFM may have poor predictions on some of the augmented entries due to false-positive protein-peptide interactions. This could lead to adding noise into the training set. Therefore, we only selected augmented entries with global backbone RMSD less than 5Å, which resulted in 1,137 augmented training entries.

To find additional data for testing, we scraped new entries from the PDB past the AF training cutoff of 2018-04-30 on 2022-12-07. We kept cases in which the biounit had a resolution of 3.0 Å or higher, only contained protein chains, and had at least two chains. One chain must have no more than 60, and no less than 10 resolved residues, and no more than 80 total residues. This chain was labeled as the peptide ligand. Another chain must be present with at least 150 resolved residues, which was labeled as the receptor. Data that shared a PDB ID with training cases already present in the PepBDB data was removed, and remaining cases were cleaned and renumbered, clustered via MMseqs2 against all prior data to remove redundancy, and created AlphaFold embeddings resulting in 837 new entries across 466 new clusters. We repeated this process on 2023-09-10 collecting data deposited since 2022-12-07 to generate a total of 46 new test cases.

### 3.2 Evaluation Metrics

To evaluate the predicted peptide-protein complex structures, we employed the traditional CAPRI docking metrics. These include the Interface RMSD (I-RMSD) which measures the mainchain RMSD of atoms at the interaction interface, Ligand RMSD (L-RMSD) which measures the mainchain RMSD of the peptide ligand when the receptors are superimposed, and the Fraction of native contacts (F_nat_) which measures how many of the native contacts are present within the predicted model. We also used the DockQ score [30] to measure the performance of our model against several baseline methods. DockQ is a composite metric of the three traditional CAPRI metrics and produces a score between 0 (worst) – 1 (best). DockQ scores larger than 0.8 are considered as ‘high’ structure quality, and less than 0.23 are considered as ‘incorrect’ structure quality.

### 3.3 Implementation Details

We implemented DistPepFold in PyTorch. All experiments were carried out on a single RTX 6000 GPU with 48 GB memory. We used OpenFold implementation of AF training code [31]. We initialized the structure module for both the teacher and student models with AFM’s structure module weights. Evoformer blocks in both the teacher and student models were randomly initialized. For both the teacher and student models, we used AdamW optimizer with a learning rate of 1e-3 and batch size of 4 with gradient accumulation steps of 8. We adopted Cosine Annealing over a total of 200 training epochs. We slightly modified the continuous cropping algorithm in AFM to ensure that the peptide residues are always included during training. The teacher model and student model are trained separately with the teacher model’s weights fixed during the student model training. During student training, we set hyperparamter α to 0.1.

## 4. Result and Discussion

A quantitative comparison of structure prediction performance is shown in **Table 1**. During evaluation, we first applied sequence-based alignment on the receptor structure, and then metrics were computed with respect to the peptide structure. To get the AFM prediction, we used AFM v2.2.0 weights with the default parameter settings as proposed previously [10]. We extracted single and pair representations from the last iteration of recycle and use these as input to our methods. We compared against AFM with several methods: 1) Structure Module: Simply finetuning the structure module. 2) Structure Module + Evoformer: same architecture as student model but without distillation loss during training. 3) Student Model: our proposed student model trained using privileged knowledge distillation. 4) Student Model + Augmentation: our proposed student model trained using privileged knowledge distillation and additional augmentation data. As shown in **Table 1**, the student model outperformed AFM across all evaluation metrics. In general, AFM achieved decent performance in protein peptide docking performance with average DockQ scores of 0.515 across 92 targets in the test set. However, we noticed that we gain slight improvement across all metrics by finetuning the structure module. This indicates that the additional finetuning with protein-peptide data is effective. Similarly, adding Evoformer blocks can further improve structure prediction performance of our model. The student model outperformed all other methods, indicating that it is able to learn useful information through the knowledge distillation process. Use of additional augmentation data did result in slight improvement in model performance, indicating that our method can benefit from more training data. However, we observed that there were more incorrect binding site predictions. This might be because too many false positive protein-peptide interactions exist within the augmentation data, which introduces too much noise during training.

**Table 1:** Peptide modeling quality for DistPepFold models and AFM. Docking quality metrics were calculated and averaged for predicted structures generated by our models, as well as AFM for all 92 targets in the test set. L-RMSD and I-RMSD scores decrease with increasing structure quality. F_nat_ and DockQ both increase with increasing structure quality and range between 0 and 1.

| Test set | L-RMSD | I-RMSD | F<sub>nat</sub> | DockQ |
| --- | --- | --- | --- | --- |
| AFM | 14.30 | 6.37 | 0.559 | 0.515 |
| Structure Module | 11.00 | 5.09 | 0.570 | 0.519 |
| Structure Module + Evoformer | 9.99 | 4.83 | 0.564 | 0.519 |
| Student Model | **8.38** | **4.34** | 0.578 | 0.529 |
| Student Model + Augmentation | 8.85 | 4.39 | **0.592** | **0.532** |

We further investigate the improvements of the student model against AFM in **Figure 2**. Across all metrics, we see that the majority of student predicted structures in the test set have similar performance to that of AFM structures. However, for a fraction of targets the student model was able to make great improvements over AFM. This trend were more prevalent in cases where AFM had significant modeling errors (as indicated by low F_nat_ and DockQ, and high I-RMSD and L-RMSD). These significant improvements indicate the student model fixing key modeling errors in binding site identification, peptide orientation, and peptide-protein contacts over AFM.

**Figure 2.**
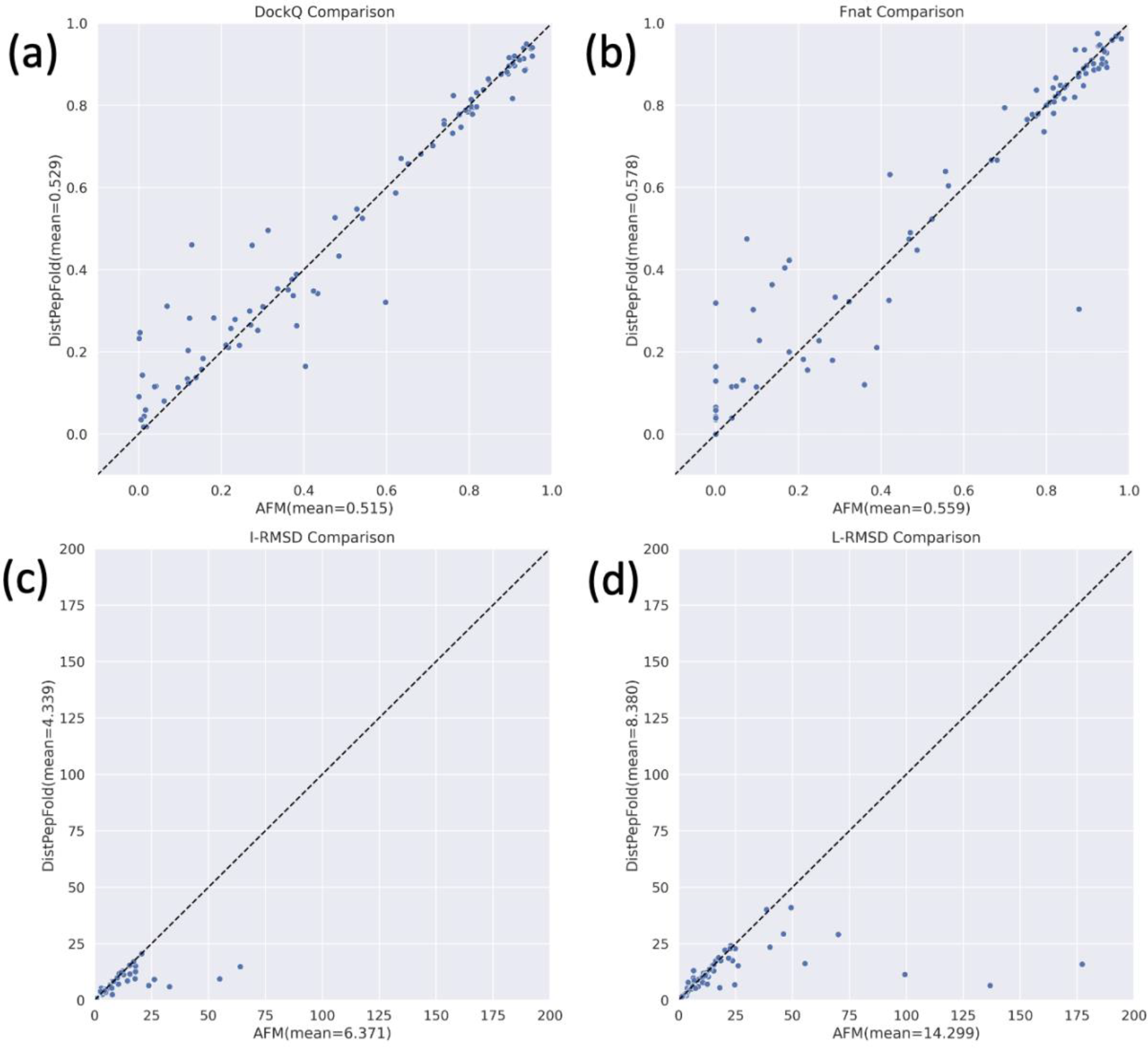
Docking metric comparison between the student model and AFM. The a) DockQ score, b) F_nat_, c) I-RMSD, and d) L-RMSD were computed and compared between the student model and AFM across all targets in the test set. The dashed lines indicate where the AFM model and student model performance would be equivalent.

In **Table 2**, we show comparison on targets in the test set with low confidence values. Here, we use AFM’s ptm + iptm score as confidence [10], which can be directly obtained from AFM’s output. We first observed that the confidence score correlates with DockQ score, where targets with high confidence tend to have high DockQ scores and targets with low confidence have low DockQ scores. There are few targets where AFM is confident but DockQ score is low. Out of 92 targets in the test set, there are 36 targets with confidence < 0.7. **Table 2** shows that the student model made clear improvements compared to AFM, with 3.1% improvements in DockQ score. This demonstrates that the student is able to make structure improvements based on AFM predicted structures. We observed that models trained without knowledge distillation, i.e. Structure Module, Structure Module + Evoformer, did not produce structure variations in the predictions compared to the student model. Instead, they tend to only make slight movements of peptide structure, which resulted in marginal improvements on F_nat_ and improved DockQ score.

**Table 2:** Peptide modeling quality for low confidence structures. For 36 targets in the test set where AFM has a low confidence score (ptm + iptm < 0.7) averages of docking quality metrics were calculated for predicted structures from AFM as well as our models. L-RMSD and I-RMSD scores decrease with increasing structure quality. F_nat_ and DockQ both increase with increasing structure quality and range between 0 and 1.

| Test set | L-RMSD | I-RMSD | $F_{\text{nat}}$ | DockQ |
| --- | --- | --- | --- | --- |
| AFM | 26.08 | 10.92 | 0.262 | 0.243 |
| Structure Module | 17.65 | 7.83 | 0.272 | 0.250 |
| Structure Module + Evoformer | 16.33 | 7.46 | 0.283 | 0.255 |
| Student Model | **13.75** | **6.69** | 0.323 | 0.274 |
| Student Model + Augmentation | 14.36 | 6.76 | **0.334** | **0.280** |

In **Table 3**, we show training evaluation comparison between the structure module, student model, and teacher model. We observe that the distillation process was successful from a training point of view. The student model was able to learn from the teacher model and correct poor quality structures produced by AFM. Such effect was not observed when comparing to models trained without distillation. We noticed that there was a difference between the student model training performance and testing performance. We suspect that student generalization ability may be limited by quantity and quality of the training data. In **Figure 3**, we computed F_nat_ for each target predicted by the model in the training set and compared against AFM predictions. **Figure 3A** shows that simple finetuning of the structure module did not provide significant improvement to models with low quality F_nat_ values. However, in both the student and teacher models (**Figure 3B** and **Figure 3C**), there was substantial improvement in targets in which AFM had low quality F_nat_ values. As expected, this trend was more pronounced in the teacher model, but it was clear that this ability has successfully been transferred to the student.

**Table 3:** Comparison between AFM, the teacher model, and the student model. For all targets within the test set, docking quality metrics were calculated and averaged for predicted structures from AFM, the student model, and teacher model.

| Model | L-RMSD | I-RMSD | $F_{\text{nat}}$ | DockQ |
| --- | --- | --- | --- | --- |
| AFM | 14.35 | 6.37 | 0.559 | 0.515 |
| Teacher | **4.72** | **2.19** | **0.748** | **0.713** |
| Student | 8.39 | 4.25 | 0.581 | 0.525 |

**Figure 3.**
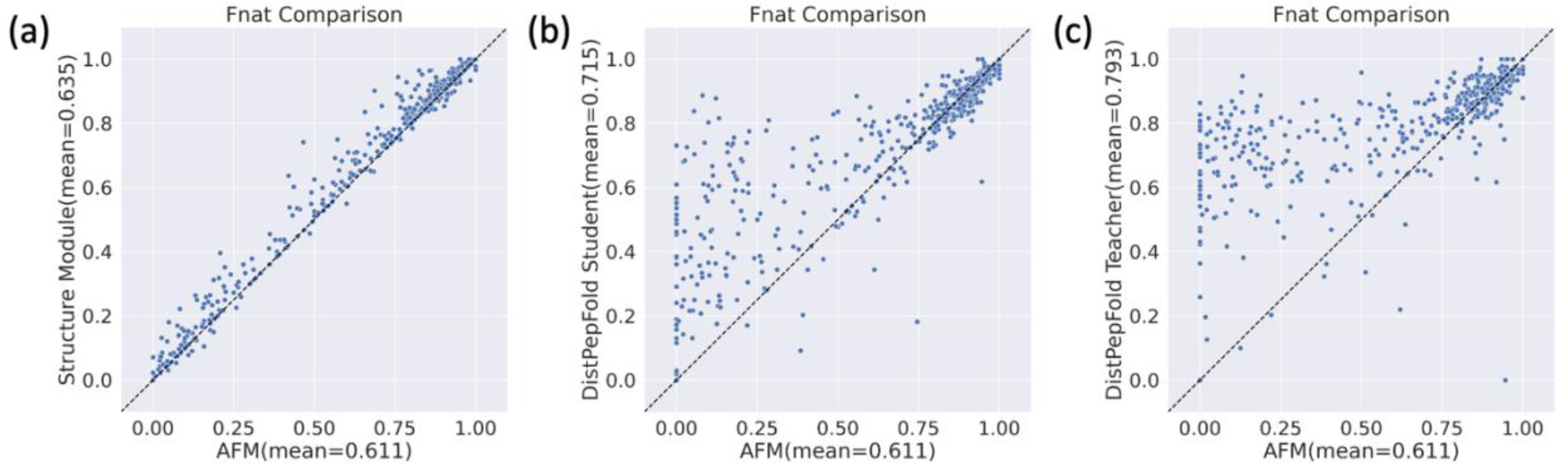
F_nat_ evaluation of the fine-tuned structure module, student model, and teacher model. F_nat_ values were calculated for all predicted structures in the training set and compared against those of AFM structures. This comparison is completed for a) finetuning of the structure module, b) the student model, and c) teacher model. The dashed lines indicate where F_nat_ values between the models would be equivalent.

In addition to AFM, we also compared DistPepFold against several traditional peptide docking methods, namely CABS-dock [32] and MDockPeP2 [33] (**Table 4**), on 46 targets that were released after the AFM training cutoff date. Unlike DistPepFold and AFM which predict the complex directly from the sequences, these traditional methods require the receptor structure, along with the peptide sequence as input and output a structure of the receptor-peptide complex structure. Among the 46 targets, we successfully run CABS-dock on and MDockPeP2 on 37 and 42 targets respectively. In addition, we reduced test redundancy by selecting the highest resolution structure within a cluster. As result, we report performance on 32 targets where we are able collect results from all method for comparison. The deep learning-based methods performed better than traditional methods by a large margin. However, we observed that the structural quality is often not satisfactory due to incorrectly identified binding sites. Our method was able to outperform all baselines across all metrics. Compared to AFM, we achieved 2.60% and 3.28% DockQ and F_nat_ improvement respectively. This is due to structural improvements on several targets and some marginal improvements on most targets. We also report performance of top-5, top-10 for CABS-dock and MdockPeP2. We selected structure with the best DockQ score among the top 10 model ranked by CABS-dock and MDockPeP2’s confidence score. We observed that both CABS-dock and MDockPeP2 benefited largely from selecting top-5, top-10 models.

**Table 4:** Comparison of DistPepFold to other methods. For 35 targets in the testing set that are not seen by AFM, we generated protein-peptide complex structures with the DistPepFold student model, AFM, CABS-Dock, and MDockPeP2, and calculated and averaged docking metrics for each method. CABS-Dock and MDockPeP2 both produce an ensemble of models, so the highest quality model within the top 10 ranked models was chosen for analysis.

| Method | L-RMSD | I-RMSD | F<sub>nat</sub> | DockQ |
| --- | --- | --- | --- | --- |
| CABS-dock | 22.90 | 8.20 | 0.092 | 0.134 |
| CABS-dock<br>(Top 5) | 14.24 | 5.25 | 0.181 | 0.233 |
| CABS-dock<br>(Top 10) | 12.31 | 4.61 | 0.213 | 0.272 |
| MDockPeP2 | 21.80 | 7.89 | 0.142 | 0.188 |
| MDockPeP2<br>(Top 5) | 14.25 | 5.08 | 0.236 | 0.290 |
| MDockPeP2<br>(Top 10) | **10.49** | **3.88** | 0.310 | 0.369 |
| AFM | 17.68 | 7.39 | 0.390 | 0.405 |
| DistPepFold | 11.27 | 5.08 | **0.428** | **0.431** |

We further investigated performance of DistPepFold against other modeling methods in terms of DockQ scores (**Figure 4**). For CABS-dock and MDockPeP2, we selected the best model ranked by score computed by the methods. The distribution of DockQ scores shows that CABS-dock and MDockPeP2 are rarely able to achieve scores above 0.4. the performance of AFM was closer to DistPepFold. However, DistPepFold was still able to make improvements on a majority of the targets over AFM, particularly those with lower DockQ scores.

**Figure 4:**
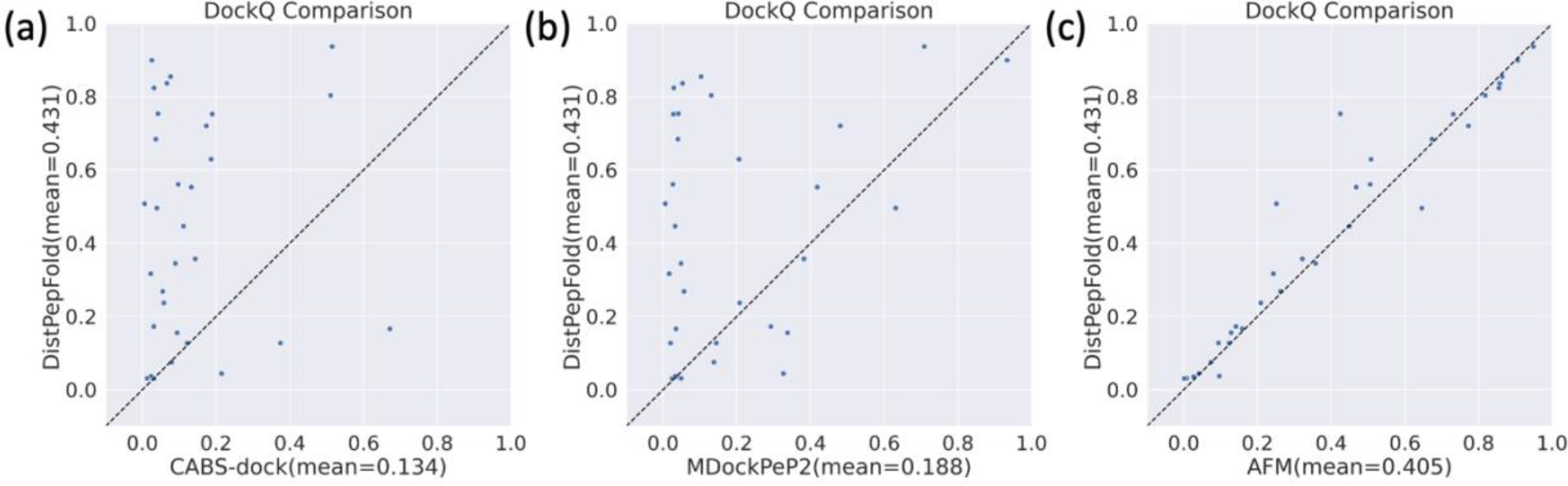
DockQ score comparison between DistPepFold and other methods. DockQ scores were calculated for structures generated by a) CABS-dock, b) MDockPeP2, and c) AFM for each of the 32 targets in the testing set that are not seen by AFM and were subsequently plotted against the scores of the DistPepFold student model. The dashed lines indicate where DockQ scores between the models would be equivalent.

To further explore how well the student learned, we inspected the pair representations before and after the student’s model. **Figure 5** shows examples of the native contact map (left) and embedding difference (right) for two targets from the test set, 3n00 and 2rqw. To get the embedding difference, we simply compute the hamming distance and normalized the value. We observe that the embedding difference tends to be higher near contact regions, particularly near peptide-protein interaction regions. This indicates that the student model wants to modify the contact region during the structure modeling process, which also increases the model’s interpretability.

**Figure 5:**
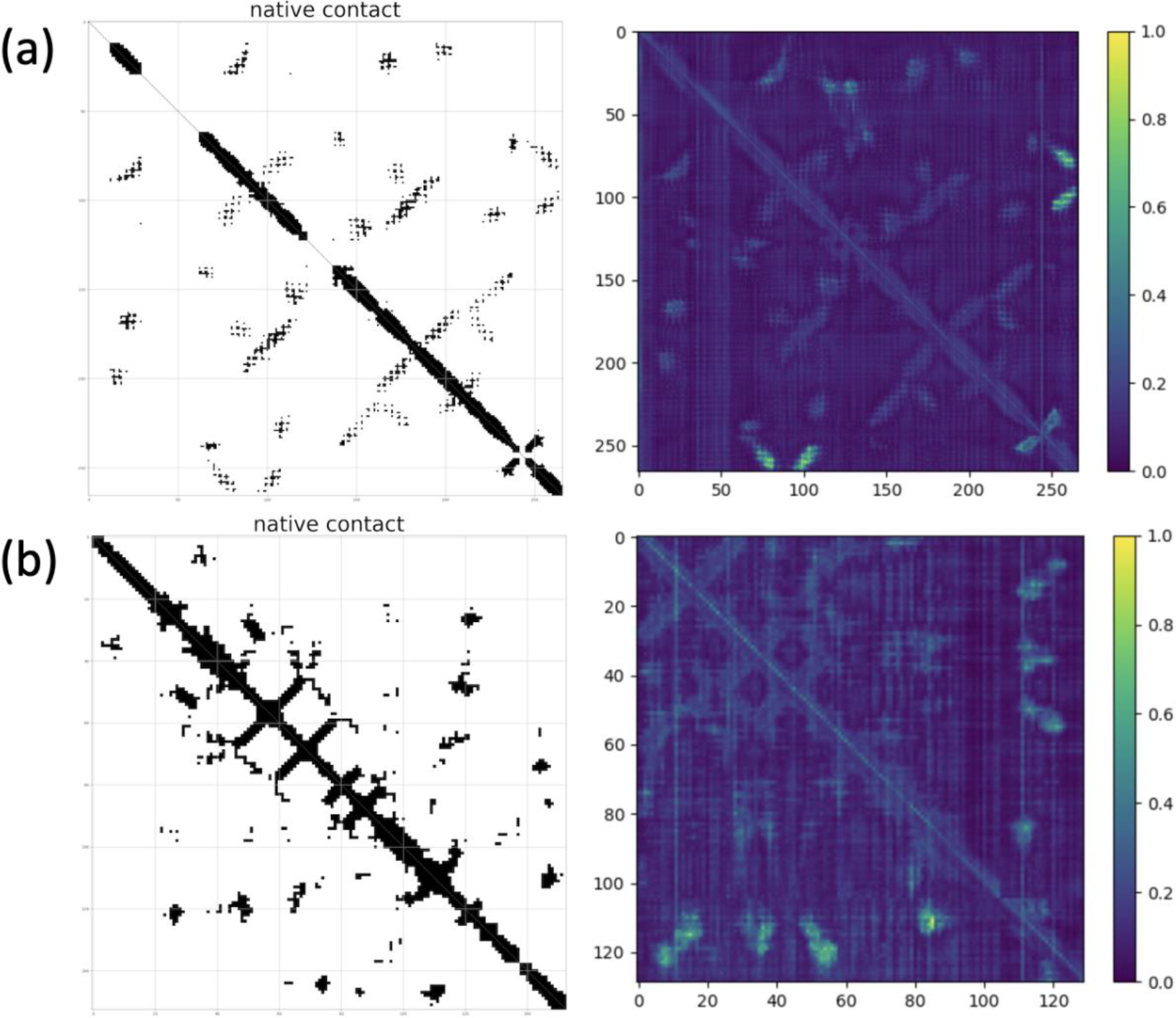
Examples of student model learning in the embedding space. The native contact map (left), and normalized embedding difference between the pair representations before and after the student model (right) are shown for two targets in the test set: 3n00 (a), and 2rqw (b). Black regions within the contact map indicate contacting residues, and brighter regions within the embedding difference indicate that the difference is large. The residue index for both the contact map and embedding difference start with the receptor sequence, and end with the peptide sequence. For 3n00 (a), the receptor spans residues 1-246, with the peptide spanning residues 247-268. For 2rqw (b), the receptor spans residues 1-106, with the peptide spanning residues 107-131.

We also analyzed the performance of the student model throughout the training process. **Figure 6** shows two examples in the validation set, 5fvd (left) and 5mlu (right). We observed that structure quality increases during the training, which indicates that student has learned from the teacher and is able to improve the structure based on AFM’s prediction. We also observed some targets are hard to optimize during the training process. This could be due to the fact that the student may have already reached its learning capability from the current dataset.

**Figure 6:**
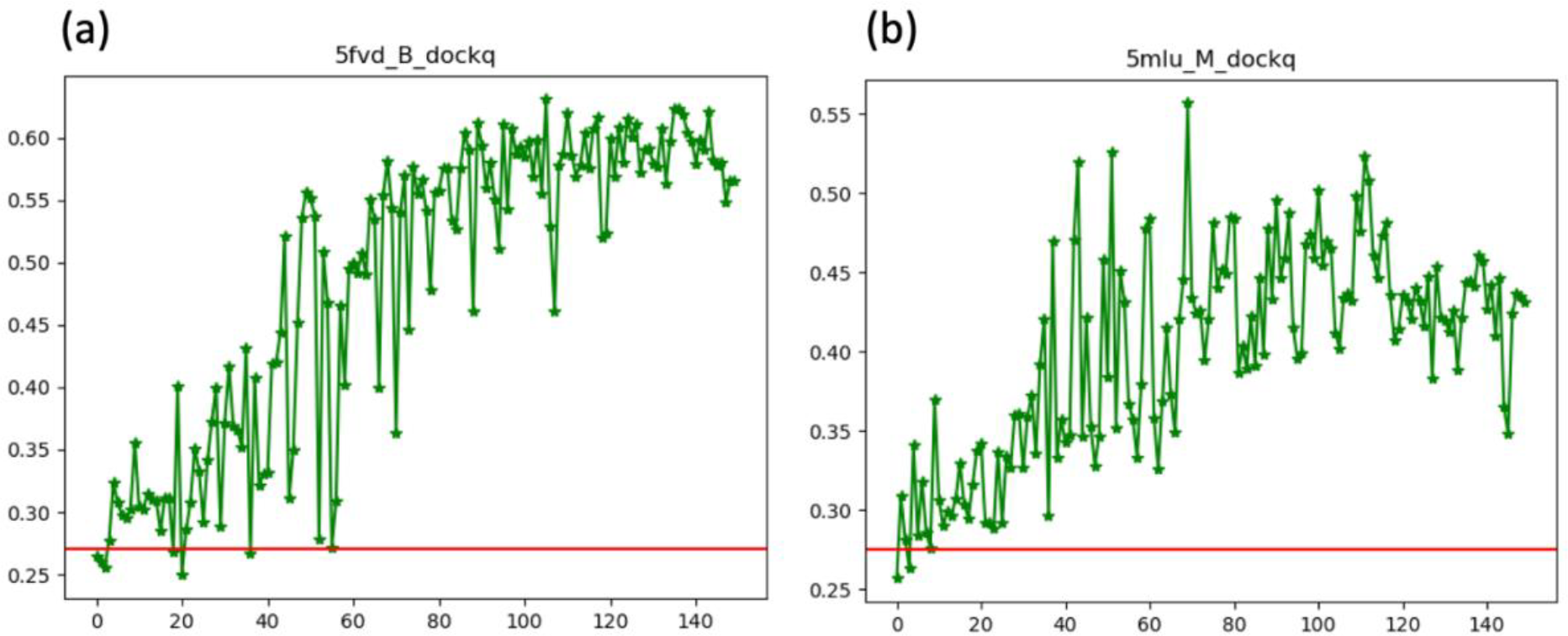
Examples of the optimization process for two validation targets: Two targets from the validation set, 5fvd (a), and 5mlu (b) had DockQ scores calculated for every epoch throughout training. The horizontal line indicates the DockQ score of AFM.

## 5. Conclusion

In this work, we proposed DistPepFold for end-to-end structure prediction of protein-peptide complexes. We developed a teacher-student framework and improved the performance of structure prediction through privileged knowledge distillation, in which the teacher has privileged knowledge regarding protein contacts that can be learned by the student model. Throughout our experiments, we showed that the teacher model is able to leverage the native contact information and predict near-native structures, and that the student model is able to learn from the teacher to make structure improvements. Benchmarking against traditional docking protocols CABS-dock and MDockPeP2 showed substantial improvement in performance, as well as moderately better performance over AFM, including several cases where DistPepFold was able to correctly identify binding sites where AFM was not. Future work on Peptide-Protein docking may address limitations in model’s generalization performance. The source code, training, and testing protocols for DistPepFold, including model weights, are available on the Kihara lab GitHub page at https://github.com/kiharalab/DistPepFold.

## Acknowledgment

This work was partly supported by the National Institutes of Health (R01GM133840) and by the National Science Foundation (DBI2003635, DBI2146026, IIS2211598, DMS2151678, CMMI1825941, and MCB1925643).

## Notes

### Competing Interest Statement

The authors have declared no competing interest.

